# *U2AF1* mutations alter splice site recognition in hematological malignancies

**DOI:** 10.1101/001107

**Authors:** Janine O. Ilagan, Aravind Ramakrishnan, Brian Hayes, Michele E. Murphy, Ahmad S. Zebari, Philip Bradley, Robert K. Bradley

## Abstract

Whole-exome sequencing studies have identified common mutations affecting genes encoding components of the RNA splicing machinery in hematological malignancies. Here, we sought to determine how mutations affecting the 3′ splice site recognition factor U2AF1 alter its normal role in RNA splicing. We find that *U2AF1* mutations influence the similarity of splicing programs in leukemias, but do not give rise to widespread splicing failure. *U2AF1* mutations cause differential splicing of hundreds of genes, affecting biological pathways such as DNA methylation (*DNMT3B*), X chromosome inactivation (*H2AFY*), the DNA damage response (*ATR, FANCA*), and apoptosis (*CASP8*). We show that *U2AF1* mutations alter the preferred 3′ splice site motif in patients, in cell culture, and *in vitro*. Mutations affecting the first and second zinc fingers give rise to different alterations in splice site preference and largely distinct downstream splicing programs. These allele-specific effects are consistent with a computationally predicted model of U2AF1 in complex with RNA. Our findings suggest that *U2AF1* mutations contribute to pathogenesis by causing quantitative changes in splicing that affect diverse cellular pathways, and give insight into the normal function of U2AF1’s zinc finger domains.

## Introduction

Myelodysplastic syndromes (MDS) represent a heterogeneous group of blood disorders characterized by dysplastic and ineffective hematopoiesis. Patients frequently suffer from cytopenias, and are at increased risk for disease transformation to acute myeloid leukemia (AML) (Tefferi and Vardiman 2009). The only curative treatment is hematopoietic stem cell transplantation, for which most patients are ineligible due to advanced age at diagnosis. The development of new therapies has been slowed by our incomplete understanding of the molecular mechanisms underlying the disease.

Recent sequencing studies of MDS patient exomes identified common mutations affecting genes encoding components of the RNA splicing machinery, with ~45-85% of patients affected (Yoshida et al. 2011; Papaemmanuil et al. 2011; Visconte et al. 2011; Graubert et al. 2011). Spliceosomal genes are the most common targets of somatic point mutations in MDS, suggesting that dysregulated splicing may constitute a common theme linking the disparate disorders that comprise MDS. Just four genes—*SF3B1, SRSF2, U2AF1*, and *ZRSR2*—carry the bulk of the mutations, which are mutually exclusive and occur in heterozygous contexts (Yoshida et al. 2011). Targeted sequencing studies identified high-frequency mutations in these genes in other hematological malignancies as well, including chronic myelomonocytic leukemia and AML with myelodysplastic features (Yoshida et al. 2011). Of the four commonly mutated genes, *SF3B1, U2AF1*, and *ZRSR2* encode proteins involved in 3′ splice site recognition (Cvitkovic and Jurica 2012; Shen et al. 2010), suggesting that altered 3′ splice site recognition is an important feature of the pathogenesis of MDS and related myeloid neoplasms.

*U2AF1* (also known as *U2AF35*) may provide a useful model system to dissect the molecular consequences of MDS-associated spliceosomal gene mutations. *U2AF1* mutations are highly specific—they uniformly affect the S34 and Q157 residues within the first and second CCCH zinc fingers of the protein—making comprehensive studies of all mutant alleles feasible (Fig. 1A). Furthermore, U2AF1’s biochemical role in binding the AG dinucleotide of the 3′ splice site is relatively well-defined (Wu et al. 1999; Zorio and Blumenthal 1999; Merendino et al. 1999). U2AF1 preferentially recognizes the core RNA sequence motif yAG|r (Fig. 1B), which matches the genomic consensus 3′ splice site and intron|exon boundary that crosslinks with U2AF1 (Wu et al. 1999). Nevertheless, our understanding of U2AF1:RNA interactions is incomplete. U2AF1’s U2AF homology motif (UHM) is known to mediate U2AF1:U2AF2 heterodimer formation (Kielkopf et al. 2001); however, both the specific protein domains that give rise to U2AF1’s RNA binding specificity and the normal function of U2AF1’s zinc fingers are unknown. Accordingly, the precise mechanistic consequences of *U2AF1* mutations are difficult to predict.

**Figure 1.**
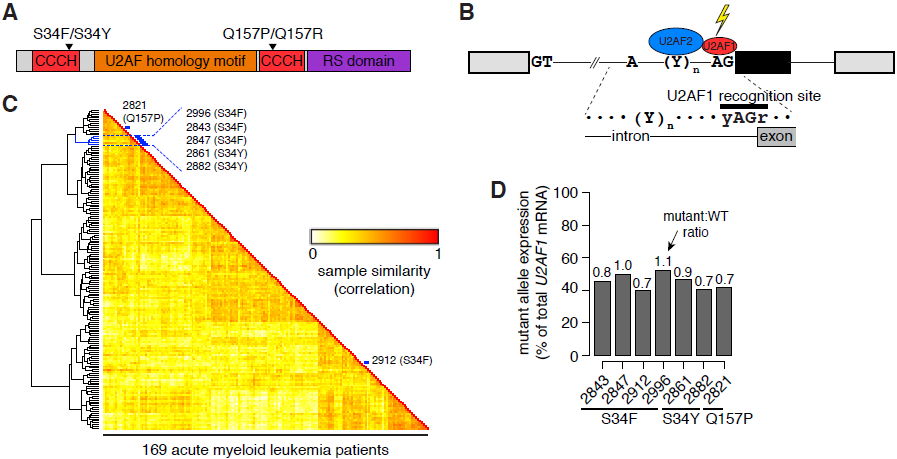
*U2AF1* mutations contribute to splicing programs in AML. (A) U2AF1 domain structure (The UniProt Consortium 2012; Kielkopf et al. 2001) and common mutations. CCCH, CCCH zinc finger. (B) Schematic of U2AF1 interaction with the 3′ splice site of a cassette exon (black). (C) Heat map illustrating similarity of alternative splicing programs in AML transcriptomes. Dendrogram is from an unsupervised cluster analysis based on cassette exon inclusion levels. Blue, samples with *U2AF1* mutations. (D) *U2AF1* mutant allele expression as a percentage of total *U2AF1* mRNA in AML transcriptomes. Numbers above bars indicate the ratio of mutant to WT allele expression.

Since the initial reports of common *U2AF1* mutations in MDS, the molecular consequences of *U2AF1* mutations have been controversial. An early study found that overexpression of mutant U2AF1 in HeLa cells resulted in dysfunctional splicing marked by frequent inclusion of premature termination codons and intron retention (Yoshida et al. 2011), while another early study reported increased exon skipping in a minigene assay following mutant U2AF1 expression in 293T cells, as well as increased cryptic splice site usage in the *FMR1* gene in MDS samples (Graubert et al. 2011). *U2AF1* mutations have been suggested to cause both alteration/gain of function (Graubert et al. 2011) and loss of function (Yoshida et al. 2011; Makishima et al. 2012). More recently, two studies analyzed acute myeloid leukemia transcriptomes from The Cancer Genome Atlas (TCGA), and found that exons with increased or decreased inclusion in samples with *U2AF1* mutations exhibited different nucleotides prior to the AG of the 3′ splice site (Przychodzen et al. 2013; Brooks et al. 2014), suggesting that *U2AF1* mutations may cause specific alterations to the RNA splicing process.

To determine how *U2AF1* mutations alter RNA splicing in hematopoietic cells, we combined patient data, cell culture experiments, and biochemical studies. We found that *U2AF1* mutations cause splicing alterations in biological pathways previously implicated in myeloid malignancies, including epigenetic regulation and the DNA damage response. *U2AF1* mutations drive differential splicing by altering the preferred 3′ splice site motif in an allele-specific manner. Our results identify downstream targets of *U2AF1* mutations that may contribute to pathogenesis, show that different *U2AF1* mutations are not functionally equivalent, and give insight into the normal function of U2AF1’s zinc finger domains.

## Results

### *U2AF1* mutations are associated with distinct splicing programs in AML

We first tested whether *U2AF1* mutations were relevant to splicing programs in leukemias with an unbiased approach. We quantified genome-wide cassette exon splicing in the transcriptomes of 169 *de novo* adult acute myeloid leukemia (AML) samples that were sequenced as part of The Cancer Genome Atlas (The Cancer Genome Atlas Research Network 2013) and performed unsupervised cluster analysis. Five of the seven samples carrying a *U2AF1* mutation clustered together (Fig. 1C). One of the samples that fell outside of this cluster had a Q157 rather than S34 *U2AF1* mutation, and the other carried a mutation in the putative RNA splicing gene *KHDRBS3* in addition to a *U2AF1* mutation, potentially contributing to its placement in an outgroup. Both of the outgroup *U2AF1* mutant samples additionally had low mutant allele expression relative to wild-type (WT) allele expression (Fig. 1D). These results suggest that *U2AF1* mutations are associated with distinct splicing patterns in patients, and are consistent with a recent report that spliceosomal mutations define a subgroup of myeloid malignancies based on gene expression and DNA methylation patterns (Taskesen et al. 2014).

### *U2AF1* mutations alter RNA splicing in blood cells

To determine how *U2AF1* mutations affect RNA splicing in an experimentally tractable system, we generated K562 erythroleukemic cell lines that stably expressed transgenic FLAG-tagged U2AF1 protein (WT, S34F, S34Y, Q157P, or Q157R mutations) at modest levels in the presence of the endogenous protein (Fig. 2A). This expression strategy, where the transgene was modestly overexpressed at levels of 1.8-4.7X endogenous *U2AF1* (Fig. 2B), is consistent with the coexpression of WT and mutant alleles at approximately equal levels that we observed in AML transcriptomes. Similar co-expression of WT and mutant alleles has been previously reported in MDS patients carrying *U2AF1* mutations (Graubert et al. 2011). We separately knocked down (KD) endogenous *U2AF1* to ~13% of normal U2AF1 protein levels in the absence of transgenic expression to test whether the mutations cause gain or loss of function (Fig. 2A).

**Figure 2.**
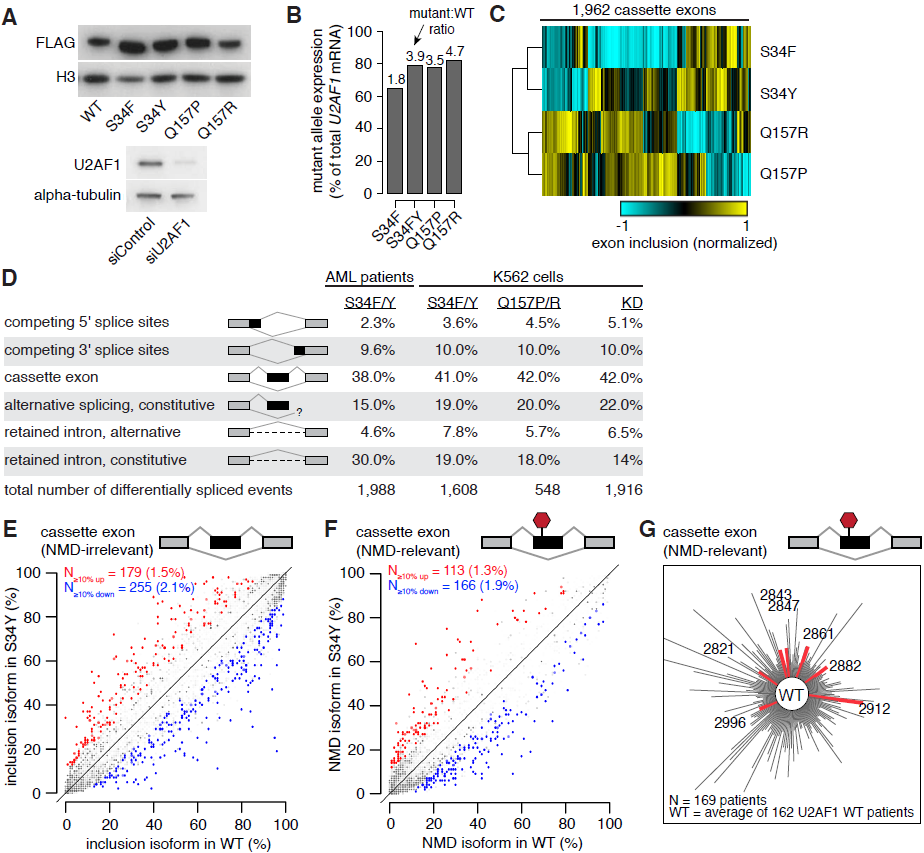
*U2AF1* mutations alter splicing, but do not cause splicing failure. (A) Western blots showing levels of FLAG-tagged U2AF1 in K562 cells stably expressing the indicated alleles (top) and levels of endogenous U2AF1 in K562 cells following transfection with a non-targeting siRNA or a siRNA pool against *U2AF1* (bottom). (B) *U2AF1* mutant allele expression as a percentage of total *U2AF1* mRNA in K562 cells. (C) Heat map of K562 cells expressing mutant *U2AF1*. Dendrogram is from an unsupervised cluster analysis based on cassette exon inclusion levels. (D) *U2AF1* mutation-dependent changes in splicing for AML S34 vs. WT patients, K562 S34F or S34Y vs. WT expression, K562 Q157P or Q157R vs. WT expression, and K562 *U2AF1* KD vs. control KD. Percentages indicate the fraction of mutation-dependent splicing changes falling into each category of splicing event. (E) Levels of cassette exon inclusion in K562 cells expressing WT or S34Y *U2AF1*. N, numbers of alternatively spliced cassette exons with increased/decreased inclusion. Percentages, fraction of alternatively spliced cassette exons that are affected by S34Y expression. Events that do not change significantly are rendered transparent. Plot restricted to cassette exon events that are predicted to not induce nonsense-mediated decay (NMD). (F)Levels of NMD-inducing isoforms of cassette exon events in K562 cells expressing WT or S34Y *U2AF1*. (G)Levels of NMD-inducing isoforms of cassette exon events in AML transcriptomes. Distance from the center measures the splicing dissimilarity between each AML transcriptome and the average of all *U2AF1* WT samples, defined as the sum of absolute differences in expression of NMD-inducing isoforms.

To identify mutation-dependent changes in splicing, we performed deep RNA-seq on these K562 cell lines stably expressing each mutant protein and on the control and *U2AF1* KD cells (~100M 2×49 bp reads per cell line). This provided sufficient read coverage to measure quantitative inclusion of ~20,000 cassette exons that were alternatively spliced in K562 cells. Unsupervised cluster analysis of global cassette exon inclusion in these cell lines placed S34F/Y and Q157P/R as distinct groups and revealed that mutations within the first and second zinc fingers are associated with largely distinct patterns of exon inclusion (Fig. 2C). This is consistent with our cluster analysis of AML transcriptomes, where the one sample with a Q157 mutation was placed as an outgroup to samples with S34 mutations.

We next assembled comprehensive maps of splicing changes driven by *U2AF1* mutations in AML transcriptomes, K562 cells expressing mutant *U2AF1*, and K562 cells following *U2AF1* KD. We tested ~125,000 annotated alternative splicing events for differential splicing, and assayed ~160,000 constitutive splice junctions for evidence of novel alternative splicing or intron retention. We required a minimum change in isoform ratio of 10% to call an event differentially spliced. As our cluster analysis of K562 cells indicated that S34 and Q157 mutations generated distinct splicing patterns, we compared the six S34 AML samples to all *U2AF1* WT AML samples. We separately identified splicing changes caused by both S34F and S34Y or both Q157P and Q157R in K562 cells relative to the WT control cells. The resulting catalogs of differentially spliced events revealed that all major classes of alternative splicing events, including cassette exons, competing splice sites, and retained introns, were affected by *U2AF1* mutations (Fig. 2D, Supplemental File S1-5). Cassette exons constituted the majority of affected splicing events, followed by alternative splicing or intron retention of splice junctions annotated as constitutive in the UCSC Genome Browser (Meyer et al. 2013).

Thousands of splicing events were affected by each *U2AF1* mutation, but the fraction of differentially spliced events was relatively low. For example, >400 frame-preserving cassette exons were differentially spliced in association with S34Y vs. WT *U2AF1* mutations; however, those >400 cassette exons constituted only ~3.6% of frame-preserving cassette exons that are alternatively spliced in K562 cells (Fig. 2E). Expression of any mutant allele caused differential splicing of 2-5% of frame-preserving cassette exons, with a bias towards exon skipping (Supplemental Fig. S1A). We did not observe increased levels of retained introns or isoforms that are predicted substrates for degradation by nonsense-mediated decay (NMD) in association with any *U2AF1* mutation. Instead, constitutive intron removal appeared slightly more efficient in cells expressing mutant versus WT *U2AF1* (Fig. 2F, Supplemental Fig. S1B-C). In contrast, we did observe increased levels of predicted NMD substrates and mRNAs with unspliced introns following *U2AF1* KD (Supplemental Fig. S1B-C). Consistent with these findings in K562 cells, AML samples carrying *U2AF1* mutations did not exhibit increased levels of NMD substrates or intron retention (Fig. 2G, Supplemental Fig. S2-4). We conclude that S34 and Q157 *U2AF1* mutations cause splicing changes affecting hundreds of exons, but do not give rise to widespread splicing failure.

These results contrast with a previous report that the *U2AF1* S34F mutation causes overproduction of mRNAs slated for degradation and genome-wide intron retention (Yoshida et al. 2011). The discrepancy between those results and our observations are likely due to differing experimental designs. This previous study acutely expressed the S34F mutation at ~50X WT levels in HeLa cells, while we stably expressed each mutant protein at 1.8-4.7X WT levels in blood cells (Fig. 1D). Maintaining a balance between WT and mutant protein expression—like that observed in AML and MDS patients—may be important to maintain efficient splicing.

### *U2AF1* mutations cause differential splicing of cancer-relevant genes

We next sought to identify downstream targets of *U2AF1* mutations that might contribute to myeloid pathogenesis. We took a conservative approach of requiring differential splicing in AML transcriptomes as well as in K562 cells to help identify disease-relevant events that are likely direct consequences of *U2AF1* mutations. We intersected differentially spliced events identified in three distinct comparisons: AML S34 vs. WT samples, K562 S34 vs. WT expression, and K562 Q157 vs. WT expression. 16.8% of AML S34-associated differential splicing was phenocopied in K562 S34 cells versus 4.6% for K562 Q157 cells, consistent with allele-specific effects of *U2AF1* mutations (Fig. 3A). The relatively low overlap of ~17% between AML and K562 S34-associated differential splicing is likely due to differences in gene expression patterns between K562 and AML cells, the modest nature of splicing changes caused by *U2AF1* mutations (such that many changes fall near the border of our statistical thresholds for differential splicing), and our stringent restriction to events that are differentially spliced in association with both S34F and S34Y mutations in K562 cells. This analysis revealed 54 splicing events that were affected by both S34 and Q157 mutations in AML transcriptomes and K562 cells. When we instead intersected genes containing differentially spliced events—not requiring that identical exons or splice sites be affected—we found a substantially increased intersection of 140 genes (Table 1).

**Figure 3.**
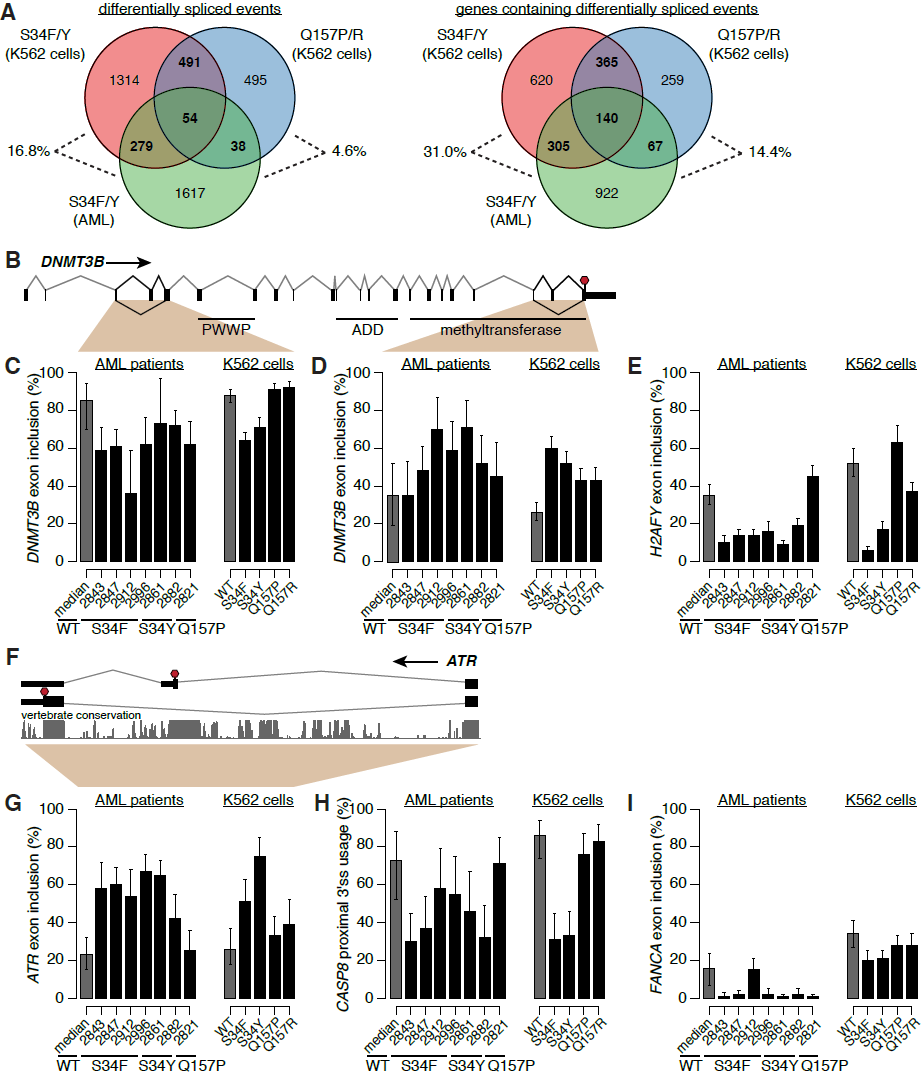
*U2AF1* mutations affect genes involved in disease-relevant cellular processes. (A) Overlap between mutation-dependent differential splicing in AML S34F/Y patients, K562 S34F/Y cells, and K562 Q157P/R cells. Overlap taken at the level of specific events (left) or genes containing differentially spliced events (right). Percentages, the fraction of differentially spliced events (left) or genes containing differentially spliced events (right) in S34F/Y AML transcriptomes that are similarly differentially spliced in K562 cells expressing S34F/Y or Q157P/R U2AF1. (B) *DNMT3B* gene structure and protein domains (The UniProt Consortium 2012). Upstream 5′ UTR not shown. PWWP, Pro-Trp-Trp-Pro domain. ADD, ATRX-DNMT3-DNMT3L domain. Red stop sign, stop codon. (C-D) Inclusion of *DNMT3B* cassette exons. Error bars, 95% confidence intervals as estimated from read coverage levels by MISO (Katz et al. 2010). (E) Inclusion of *H2AFY* cassette exon. (F) Cassette exon at 3′ end of *ATR*. Conservation is phastCons (Siepel et al. 2005) track from UCSC (Meyer et al. 2013). (G) Inclusion of cassette exon in *ATR*. (H) Usage of intron-proximal 3′ splice site of *CASP8*. (I) Inclusion of cassette exon in *FANCA*.

**Table 1.**
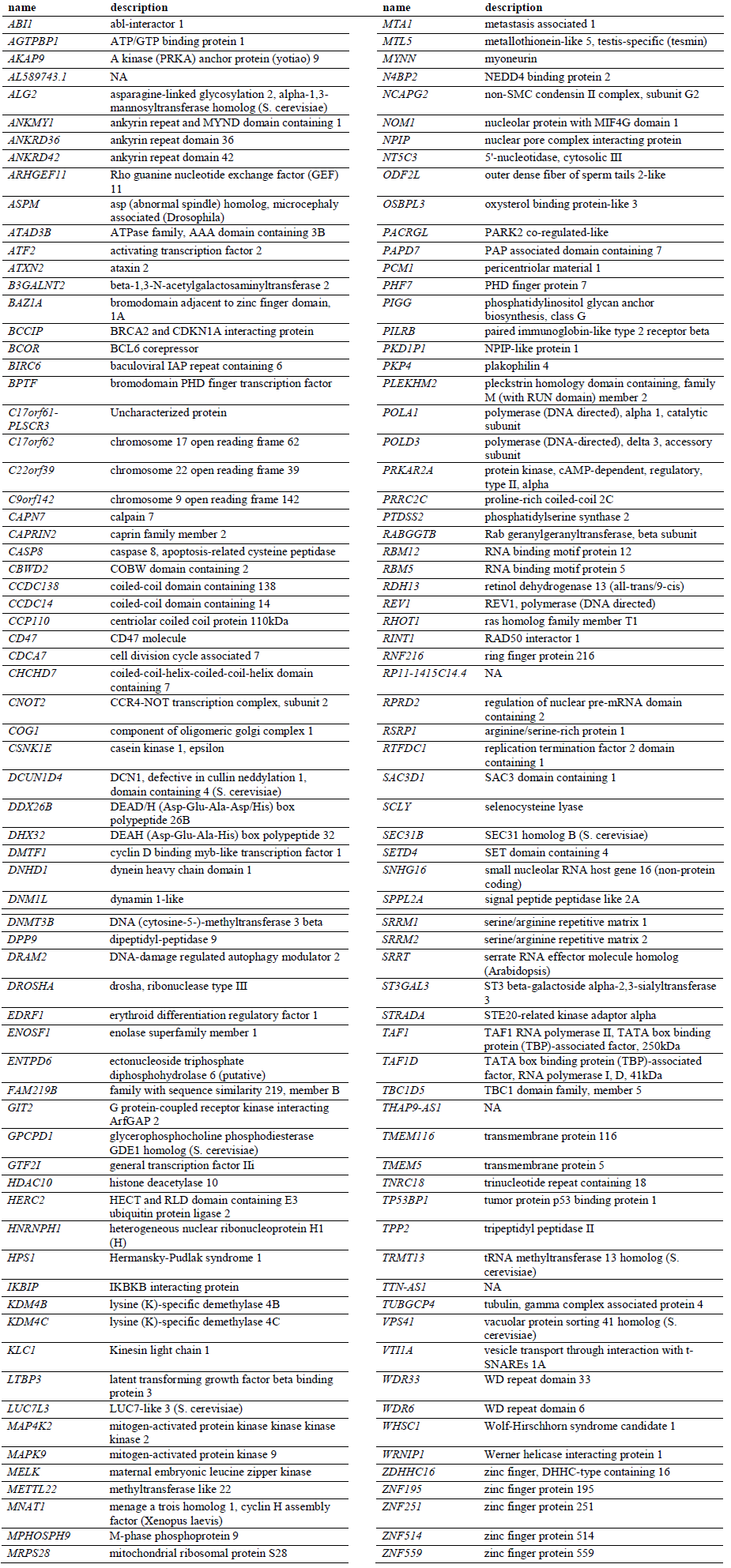
Genes that are differentially spliced in association with *U2AF1* mutations. Genes that contain events that are differentially spliced in the AML S34 samples (versus WT samples), K562 S34 samples (versus WT), and K562 Q157 samples (versus WT). Descriptions taken from Ensembl.

Many genes that were differentially spliced in association with *U2AF1* mutations participate in biological pathways previously implicated in myeloid malignancies. For example, *DNMT3A* encodes a *de novo* DNA methyltransferase and is a common mutational target in myelodysplastic syndromes and acute myeloid leukemia (Walter et al. 2011; Ley et al. 2010). Multiple exons of its paralog *DNMT3B*, including an exon encoding part of the methyltransferase domain, are differentially spliced in AML patients carrying *U2AF1* mutations as well as in K562 cells expressing *U2AF1* mutant protein (Fig. 3B–D, Supplemental Fig. S5A-B). Similarly, different exons of *ASXL1* are alternatively spliced in association with S34 mutations in AML transcriptomes and K562 cells, although the same exons are not consistently affected (Supplemental File S1-5). *ASXL1* is a common mutational target in myelodysplastic syndromes and related disorders (Gelsi-Boyer et al. 2009), and *U2AF1* and *ASXL1* mutations co-occur more frequently than expected by chance (Thol et al. 2012). Other genes participating in epigenetic processes are differentially spliced as well, such as *H2AFY* (Fig. 3E, Supplemental Fig. S5C). *H2AFY* encodes the core histone macro-H2A.1, which is important for X chromosome inactivation (Hernández-Muñoz et al. 2005). As loss of X chromosome inactivation causes a MDS-like disease in mice (Yildirim et al. 2013), differential splicing of macro-H2A.1 could potentially be relevant to disease processes.

Isoform switches, wherein a previously minor isoform becomes the major isoform, were relatively rare but did occur. For example, a cassette exon at the 3′ end of *ATR* gene, which encodes a PI3K-related kinase that activates the DNA damage checkpoint, is included at high rates in association with S34, but not Q157, mutations. This cassette exon alters the C terminus of the ATR protein, may render the mRNA susceptible to nonsense-mediated decay, and is highly conserved (Fig. 3F-G, Supplemental Fig. S5D). S34 mutations similarly cause an isoform switch from an intron-proximal to an intron-distal 3′ splice site of *CASP8* that is predicted to shorten the N terminus of the protein (Fig. 3H, Supplemental Fig. S5E).

We noticed that splicing changes frequently affected multiple genes relevant to a specific biological process, such as DNA damage (*ATR* and *FANCA*; Fig. 3G,3I, Supplemental Fig. S5D, S5F). Consistent with this observation, gene ontology analysis indicated that genes involved in the cell cycle, chromatin modification, DNA methylation, DNA repair, and RNA processing pathways, among others, are enriched for differential splicing in both AML transcriptomes and K562 cells in association with *U2AF1* mutations. This enrichment could be due to high basal rates of alternative splicing within these genes, which frequently are composed of many exons, or instead caused by specific targeting by mutant *U2AF1*. Upon correcting for gene-specific variations in the number of possible alternatively spliced isoforms, these pathways were no longer enriched in gene ontology analyses. We conclude that *U2AF1* mutations preferentially affect specific biological pathways, but that this enrichment is due to frequent alternative splicing within such genes rather than specific targeting by *U2AF1* mutant protein.

### *U2AF1* mutations cause allele-specific alterations in the 3′ splice site consensus

Previous biochemical studies showed that U2AF1 recognizes the core sequence motif yAG|r of the 3′ splice site (Wu et al. 1999; Zorio and Blumenthal 1999; Merendino et al. 1999). Accordingly, we hypothesized that the splicing changes caused by *U2AF1* mutations might be due to preferential activation or repression of 3′ splice sites in a sequence-specific manner. To test this hypothesis, we identified consensus 3′ splice sites of cassette exons that were promoted or repressed in AML transcriptomes carrying *U2AF1* mutations relative to WT patients. For each mutant *U2AF1* sample, we enumerated all cassette exons that were differentially spliced between the sample and an average *U2AF1* WT sample, requiring a minimum change in isoform ratio of 10%. Exons whose inclusion was increased or decreased in *U2AF1* mutant samples exhibited different consensus nucleotides at the −3 and +1 positions flanking the AG of the 3′ splice site. As these positions correspond to the yAG|r motif bound by *U2AF1*, this data supports our hypothesis that *U2AF1* mutations alter 3′ splice site recognition activity in a sequence-specific manner (Fig. 4A).

**Figure 4.**
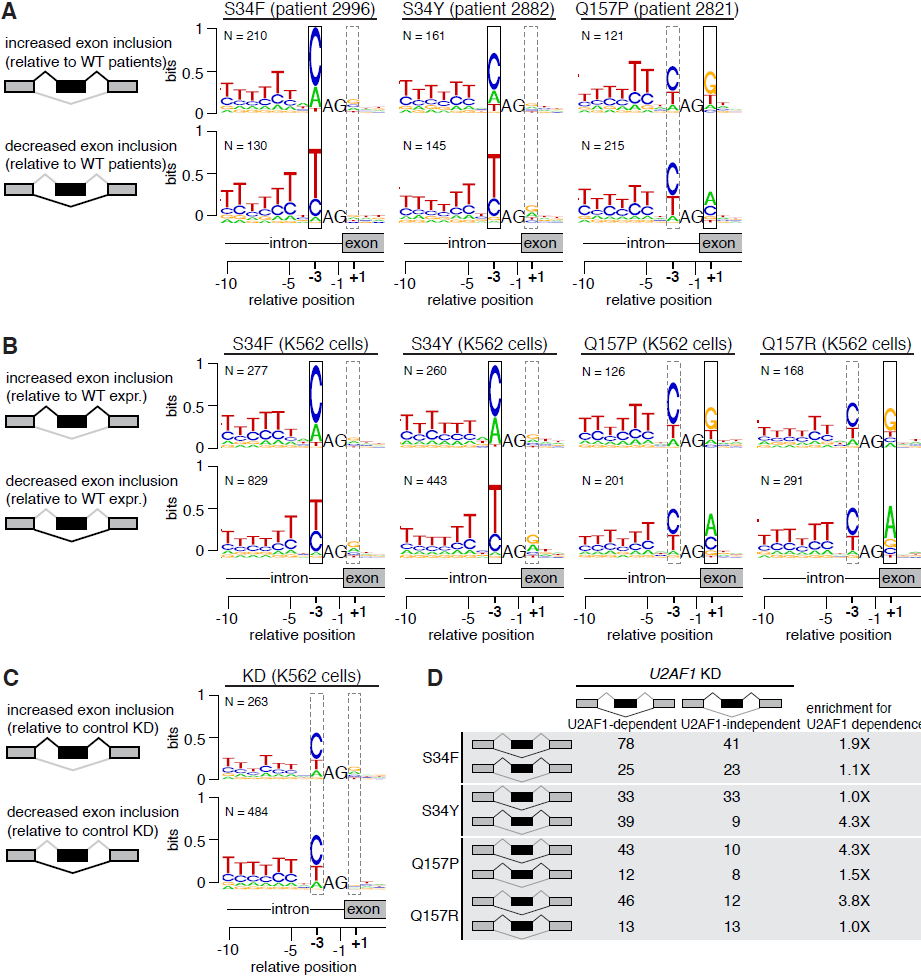
*U2AF1* mutations alter 3′ splice site consensus sequences. (A) Consensus 3′ splice sites of cassette exons with increased or decreased inclusion in *U2AF1* mutant relative to WT AML transcriptomes. Boxes highlight sequence preferences at the −3 and + 1 positions that differ from the normal 3′ splice site consensus. Vertical axis, information content in bits. N, number of cassette exons with increased or decreased inclusion in each sample. Data for all *U2AF1* mutant samples shown in Supplemental Figure S6. (B) As (A), but for K562 cells expressing the indicated mutation vs. WT. (C) As (A), but for K562 cells following *U2AF1* KD or control KD. (D) Overlap between cassette exons that are promoted or repressed by mutant vs. WT expression (rows) and *U2AF1* vs. control KD (columns) in K562 cells. Third column indicates the enrichment for U2AF1 dependence, defined as the overlap between exons affected by mutant U2AF1 expression and exons repressed versus promoted by *U2AF1* KD.

Mutations affecting different residues of U2AF1 were associated with distinct alterations in the consensus 3′ splice site motif yAG|r of differentially spliced exons. The S34F and S34Y mutations, affecting the first zinc finger, were associated with nearly identical alterations at the −3 position in all six S34 mutant samples, while the Q157P mutation, affecting the second zinc finger, was associated with alterations at the +1 position (Supplemental Fig. S6). In contrast, cassette exons that were differentially spliced in randomly chosen *U2AF1* WT samples relative to an average AML sample did not exhibit altered consensus sequences at the −3 or +1 positions (Fig. S7). These results confirm the findings of two recent studies of this cohort of AML patients—which reported a frequent preference for C instead of T at the −3 position of differentially spliced cassette exons in *U2AF1* mutant samples (Przychodzen et al. 2013; Brooks et al. 2014)—and extend their observations of altered splice site preference to show allele-specific effects of *U2AF1* mutations, which have not been previously identified.

*U2AF1* mutation-dependent sequence preferences (C/A ≫ T at the −3 position for S34F/Y and G ≫ A at the +1 position for Q157P) differ from the genomic consensus for cassette exons. C/T and G/A appear at similar frequencies at the −3 and +1 positions of 3′ splice sites of cassette exons (Supplemental Fig. S6-7), and minigene and genomic studies of competing 3′ splice sites indicate that C and T are approximately equally effective at the −3 position (Smith et al. 1993; Bradley et al. 2012). The consensus 3′ splice sites associated with promoted/repressed cassette exons in *U2AF1* mutant transcriptomes also differ from U2AF1’s known RNA binding specificity. A previous study reported a core tAG|g motif in the majority of RNA sequences bound by U2AF1 in a SELEX experiment (Wu et al. 1999). Comparing that motif with preferences observed in *U2AF1* mutant transcriptomes, we hypothesize that S34 U2AF1 promotes unusual recognition of C instead of T at the −3 position, while Q157 U2AF1 reinforces preferential recognition of G instead of A at the +1 position.

We next tested whether these alterations in 3′ splice site preference are a direct consequence of *U2AF1* mutations. Comparing K562 cells expressing mutant vs. WT *U2AF1*, we found that cassette exons that were promoted or repressed by each mutation exhibited sequence preferences at the −3 and +1 positions that were highly similar to those observed in AML patient samples (Fig. 4B). Mutations affecting identical residues (S34F/Y and Q157P/R) caused similar alterations in 3′ splice site preference, while mutations affecting different residues did not, confirming the allele-specific consequences of *U2AF1* mutations. In contrast, cassette exons that were differentially spliced following KD of endogenous *U2AF1* did not exhibit sequence-specific changes at the −3 or +1 positions of the 3′ splice site (Fig. 4C). We therefore conclude that S34 and Q157 mutations cause alteration or gain of function, consistent with the empirical absence of inactivating (nonsense or frameshift) *U2AF1* mutations observed in patients.

### *U2AF1* mutations preferentially affect U2AF1-dependent 3′ splice sites

*U2AF1* mutations are associated with altered 3′ splice site consensus sequences, yet only a relatively small fraction of cassette exons are affected by expression of *U2AF1* mutant protein. Previous biochemical studies found that only a subset of exons have “AG-dependent” 3′ splice sites that require U2AF1 binding for proper splice site recognition (Reed 1989; Wu et al. 1999). We therefore speculated that exons that are sensitive to *U2AF1* mutations might also rely upon U2AF1 recruitment for normal splicing. We empirically defined U2AF1-dependent exons as those with decreased inclusion following *U2AF1* KD, and computed the overlap between U2AF1-dependent exons and exons that were affected by *U2AF1* mutant protein expression. For every mutation, we observed an enrichment for overlap with U2AF1-dependent exons, suggesting that *U2AF1* mutations preferentially affect exons with AG-dependent splice sites (Fig. 4D).

### *U2AF1* mutations alter the preferred 3′ splice site motif yAG|r

Our genomics data shows that cassette exons promoted/repressed by *U2AF1* mutations have 3′ splice sites differing from the consensus. We therefore tested whether altering the core 3′ splice site motif of an exon influenced its recognition in the presence of WT versus mutant U2AF1. We created a minigene encoding a cassette exon of *ATR*, which responds robustly to S34 mutations in AML transcriptomes and K562 cells (Fig. 3F-G), by cloning the genomic locus containing the *ATR* cassette exon and flanking constitutive introns and exons into a plasmid. The minigene exhibited mutation-dependent splicing of the cassette exon, as expected, although cassette exon recognition was less efficient than from the endogenous locus. We then mutated the −3 position of the cassette exon’s 3′ splice site to A/C/G/T and measured cassette exon inclusion in WT and S34Y K562 cells. Robust mutation-dependent increases in splicing required the A at the −3 position found in the endogenous locus, consistent with the unusual preference for A observed in our analyses of AML and K562 transcriptomes. We additionally observed a small but reproducible increase for C (Fig. 5A). We next performed similar experiments for Q157-dependent splicing changes. We created a minigene encoding a cassette exon of *EPB49* (encoding the erythrocyte membrane protein band 4.9), mutated the +1 position of the 3′ splice site to A/C/G/T, and measured cassette exon inclusion in WT and Q157R K562 cells. Cassette exon recognition was suppressed by Q157R expression when the +1 position was an A, consistent with our genomic prediction, and was not affected by Q157R when the +1 position was mutated to another nucleotide. Therefore, for both *ATR* and *EPB49*, robust S34 and Q157-dependent changes in splicing required the endogenous nucleotides at the −3 and +1 positions.

**Figure 5.**
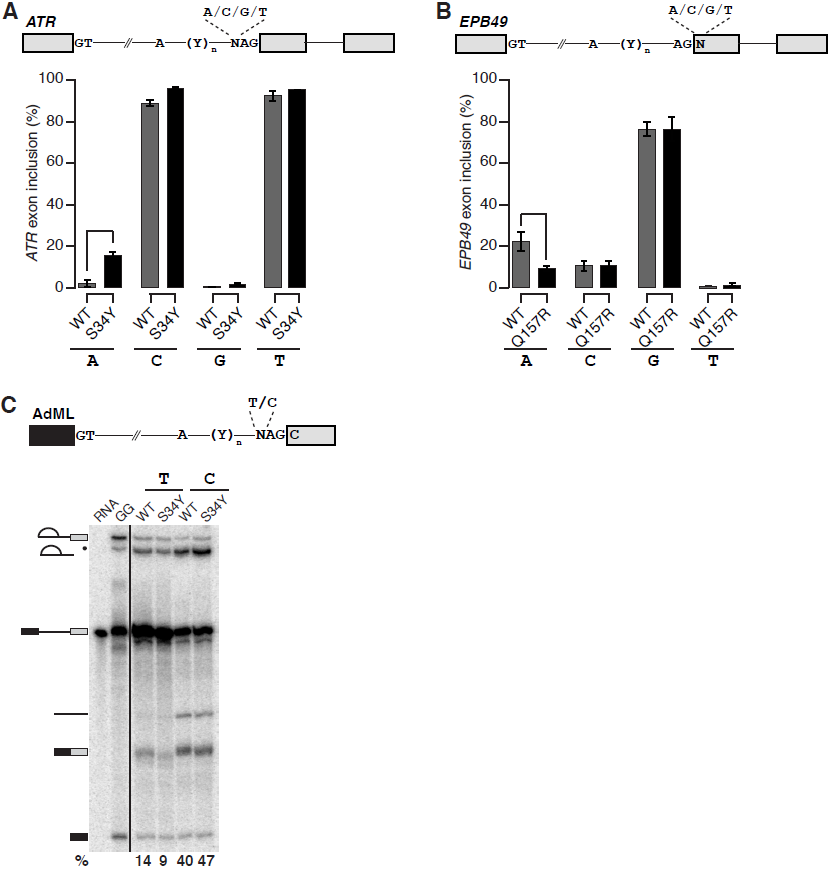
*U2AF1* mutations cause sequence-dependent changes in 3′ splice site recognition. (A) Schematic of *ATR* minigene (top) and inclusion of *ATR* cassette exon transcribed from minigenes with A/C/G/T at the −3 position of the 3′ splice site in K562 cells expressing WT or S34Y *U2AF1* (bottom). Error bars, standard deviation from biological triplicates. (B) Schematic of *EPB49* minigene (top) and inclusion of *EPB49* cassette exon transcribed from minigenes with A/C/G/T at the +1 position of the 3′ splice site in K562 cells expressing WT or Q157R *U2AF1* (bottom). (C) Schematic of AdML pre-mRNA substrate used for *in vitro* splicing (top) and *in vitro* splicing of AdML substrate incubated with nuclear extract from K562 cells expressing WT or S34Y *U2AF1* (bottom). Percentages are the fraction of second step products (spliced mRNA and lariat intron) relative to all RNA species after 60 minutes of incubation. RNA, input radiolabeled RNA. GG, pre-mRNA with the AG dinucleotide replaced by GG to illustrate the first step product of splicing. Black dot, exonucleolytic “chew back” product of the lariat intermediate.

We next tested how *U2AF1* mutations influence constitutive, rather than alternative, splicing in an *in vitro* context. We used the adenovirus major late (AdML) substrate, a standard model of constitutive splicing, and mutated the −3 position of the 3′ splice site to C/T. We measured AdML splicing efficiency following *in vitro* transcription and incubation with nuclear extract of K562 cells expressing WT or S34Y U2AF1. The AdML substrate exhibited sequence-specific changes in splicing efficiency in association with *U2AF1* mutations. Consistent with our genomic analyses, AdML with C/T at the −3 position was more/less efficiently spliced in S34Y versus WT cells (Fig. 5C). Taken together, our data demonstrate that *U2AF1* mutations cause sequence-specific alterations in the preferred 3′ splice site motif in patients, in cell culture, and *in vitro*.

### *U2AF1* mutations may modify U2AF1:RNA interactions

As *U2AF1* mutations alter the preferred 3′ splice site motif yAG|r—the same motif that is recognized and bound by U2AF1 (Wu et al. 1999)—we next investigated whether *U2AF1* mutations could potentially modify U2AF1’s RNA binding activity. U2AF1’s RNA binding specificity could originate from its U2AF homology motif (UHM) and/or its two CCCH zinc fingers. The UHM domain mediates U2AF heterodimer formation and is sufficient to promote splicing of an AG-dependent pre-mRNA substrate (Guth et al. 2001; Kielkopf et al. 2001). However, this domain binds a consensus 3′ splice site sequence with low affinity (Kielkopf et al. 2001), suggesting that it may be insufficient to generate U2AF1’s sequence specificity. As U2AF1’s zinc fingers are independently required for U2AF RNA binding (Webb and Wise 2004), and our data indicates that zinc finger mutations alter splice site preferences, we hypothesized that U2AF1’s zinc fingers might directly interact with the 3′ splice site.

To evaluate whether this hypothesis is sterically possible, we started from the experimentally determined structure of the UHM domain in complex with a peptide from U2AF2 (Kielkopf et al. 2001), modeled the conformations of the zinc finger domains bound to RNA by aligning them to the CCCH zinc finger domains in the TIS11d:RNA complex structure (Hudson et al. 2004), and sampled the conformations of the two short linker regions using fragment assembly techniques (Leaver-Fay et al. 2011). The RNA was built in two segments taken from the TIS11d complex, one anchored in the N-terminal zinc finger and one in the C-terminal finger. We modeled multiple 3′ splice site sequences (primarily variants of uuAG|ruu), and explored a range of possible alignments of the 3′ splice site within the complex. The final register was selected on the basis of energetic analysis and manual inspection using known features of the specificity pattern of the 3′ splice site (in particular, the lack of a significant genomic consensus at the −4 and +3 positions, consistent with the experimental absence of a crosslink between U2AF1 and the −4 position (Wu et al. 1999)).

Based on these simulations, we propose a theoretical model of U2AF1 in complex with RNA wherein the zinc finger domains guide recognition of the yAG|r motif, consistent with the predictions of our mutational data. The model has the following features (Fig. 6A, Supplemental File S6). The first zinc finger contacts the bases immediately preceding the splice site, including the AG dinucleotide (Fig. 6B-C), while the second zinc finger binds immediately downstream (Fig. 6D). The RNA is kinked at the splice site and bent overall throughout the complex so that both the 5′ and 3′ ends of the motif are oriented toward the UHM domain and U2AF2 peptide. Contacts compatible with the 3′ splice site consensus are observed at the sequence-constrained RNA positions. The mutated positions S34 and Q157 are nearby the bases at which perturbed splice site preferences are observed for their respective mutations. Moreover, the modified preferences can, to some extent, be rationalized by contacts seen in our simulations. S34 forms a hydrogen bond with U(-1), and preference for U at -1 appears to decrease upon mutation; the Q157P mutation would improve electrostatic complementarity with G at +1 by removing a backbone NH group, in agreement with increased G preference in this mutant.

**Figure 6.**
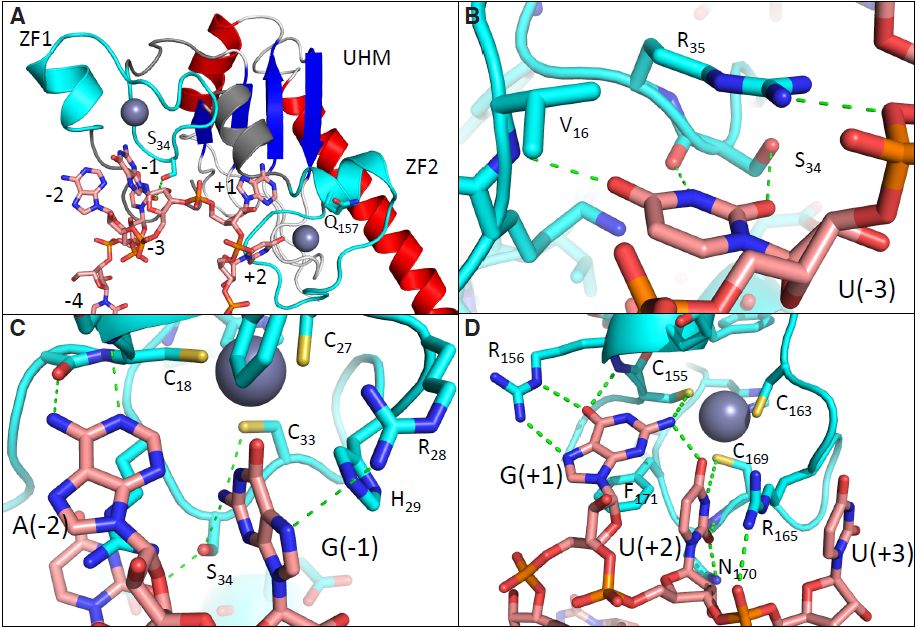
Theoretical model of the U2AF1:RNA complex. (A) Overview, with the zinc finger domains colored cyan, the RNA in salmon, and the UHM beta sheet in blue and alpha helices in red. The frequently mutated positions S34 and Q157 are shown in stick representation. ZF, zinc finger. (B-D) Interactions with individual bases characteristic of the 3′ splice site consensus. Green dotted lines indicate hydrogen bonds and favorable electrostatic interactions; RNA and selected side chains are shown in stick representation.

## Discussion

Here, we have described the mechanistic consequences of *U2AF1* mutations in hematopoietic cells, as well as provided a catalog of splicing changes driven by each common *U2AF1* mutation. *U2AF1* mutations cause highly specific alterations in 3′ splice site recognition in myeloid neoplasms. Taken together with the high frequency of mutations targeting *U2AF1* and genes encoding other 3′ splice site recognition factors, our results support the hypothesis that specific alterations in 3′ splice site recognition are important contributors to the molecular pathology of MDS and related hematological disorders.

We observed consistent differential splicing of multiple genes such as *DNMT3B* and *FANCA* that participate in molecular pathways previously implicated in blood disease. It is tempting to speculate that differential splicing of a few such genes in well-characterized pathways explain how *U2AF1* mutations drive disease. However, we instead hypothesize that spliceosomal mutations contribute to dysplastic hematopoiesis and tumorigenesis by dysregulating a multitude of genes involved in many aspects of cell physiology. This hypothesis is consistent with two notable features of our data. First, hundreds of exons are differentially spliced in response to *U2AF1* mutations. Second, many of the splicing changes are relatively modest. In both the AML and K562 data, we observed relatively few isoform switches, with the *ATR* and *CASP8* examples illustrated in Figure 3 being notable exceptions. Therefore, we expect that specific targets such as *DNMT3B* probably contribute to, but do not wholly explain, *U2AF1* mutation-induced pathophysiology. As additional data from tumor transcriptome sequencing become available—for example, as more patient transcriptomes carrying Q157 mutations are sequenced—precisely identifying disease-relevant changes in splicing will become increasingly reliable.

Our understanding of the molecular consequences of *U2AF1* mutations will also benefit from further experiments conducted during the differentiation process. Both the AML and K562 data arose from relatively “static” systems, in the sense that the bulk of the assayed cells were not actively undergoing lineage specification. *U2AF1* mutations likely cause similar changes in splice site recognition in both precursor and more differentiated cells, but altered splice site recognition could have additional consequences in specific cell types. A recent study reported that regulated intron retention is important for granulopoiesis (Wong et al. 2013), consistent with the idea that as-yet-unrecognized shifts in RNA processing may occur during hematopoiesis. By disrupting such global processes, altered splice site recognition could contribute to the ineffective hematopoiesis that characterizes MDS.

## Relevance to future studies of spliceosomal mutations

Both mechanistic and phenotypic studies of cancer-associated somatic mutations frequently focus on single mutations, even when multiple distinct mutations affecting that gene occur at high rates. Similarly, distinct mutations affecting the same gene are frequently grouped together in prognostic and other clinical studies, thereby implicitly assuming that different mutations have similar physiological consequences. Our finding that different *U2AF1* mutations are not mechanistically equivalent illustrates the value of studying all high-frequency mutations when feasible. The distinctiveness of S34 and Q157 mutation-induced alterations in 3′ splice site preference suggests that they could theoretically constitute clinically relevant disease subtypes, potentially contributing to the heterogeneity of MDS. Mutations affecting other spliceosomal genes may likewise have allele-specific consequences. For example, mutations at codons 625 versus 700 of *SF3B1* are most commonly associated with uveal melanoma (Harbour et al. 2013; Martin et al. 2013) versus MDS (Yoshida et al. 2011; Graubert et al. 2011; Papaemmanuil et al. 2011; Visconte et al. 2011) and chronic lymphocytic leukemia (Quesada et al. 2011). Accordingly, we speculate that stratifying patients by mutation could prove fruitful for future studies of spliceosomal gene mutations.

Our study additionally illustrates how investigating disease-associated somatic mutations can give insight into the normal function of proteins. With a fairly restricted set of assumptions, we computationally predicted a family of models in which the first zinc finger of U2AF1 recognizes the AG dinucleotide of the 3′ splice site. As a computational prediction, the model must be tested with future experiments. Nonetheless, given the concordance between our theoretical model of U2AF1:RNA interactions and our mutational data, this model may provide a useful framework for future studies of U2AF1 function in both healthy and diseased cells.

## Methods

### Vector construction and cell culture

Inserts encoding bicistronic constructs of the form U2AF1 + Gly Gly + FLAG + T2A + mCherry were created by standard methods (details in Supplemental Methods). These inserts were cloned into the self-inactivating lentiviral vector pRRLSIN.cPPT.PGK-GFP.WPRE (Addgene Plasmid 12252). The resulting plasmids co-express U2AF1 and mCherry under control of the PGK promoter. K562 erythroleukemia cells were grown in RPMI-1640 supplemented with 10% FCS. To generate stable cell lines, K562 cells were infected with concentrated lentiviral supernatants at a MOI of ~5 in growth media supplemented with 8 ug/mL protamine sulfate. Cells were then expanded and transduced cells expressing mCherry were isolated by fluorescence activated cell sorting (FACS) using a Becton Dickinson FACSAria II equipped with a 561 nm laser. For RNAi studies, K562 cells were transfected with a control (non-targeting) siRNA (Dharmacon D-001810-03-20) or a siRNA pool against *U2AF1* (Dharmacon ON-TARGETplus SMARTpool L-012325-01-0005) using the Nucleofector II device from Lonza with the Cell Line Nucleofector Kit V (program T16), and RNA and protein were collected 48 hours after transfection.

### mRNA sequencing

Total RNA was obtained by lysing 10 million K562 cells for each sample in TRIzol and RNA was extracted using Qiagen RNeasy columns. Using 4 ug of total RNA, we prepared poly(A)-selected, unstranded libraries for Illumina sequencing using a modified version of the TruSeq protocol (details in Supplemental Methods). RNA-seq libraries were then sequenced on the Illumina HiSeq 2000 to a depth of ~100 million 2×49 bp reads per sample.

### Accession numbers

For the AML analysis, BAM files were downloaded from CGHub (“LAML” project) and converted to FASTQ files of unaligned reads for subsequent read mapping. For the HeLa cell analysis, FASTQ files were downloaded from DDBJ series DRA000503 (http://trace.ddbj.nig.ac.jp/DRASearch/), and the reads were trimmed to 50 bp (after removing the first five bp) to restrict to the high-quality portion of the reads. A similar trimming procedure was performed in the original manuscript (Yoshida et al. 2011).

### Genome annotations and read mapping

MISO v2.0 annotations were used for cassette exon, competing 5′ and 3′ splice sites, and retained intron events (Katz et al. 2010). Constitutive junctions were defined as splice junctions that were not alternatively spliced in any isoform of the UCSC knownGene track (Meyer et al. 2013). For read mapping purposes, a gene annotation file was created by combining isoforms from the MISO v2.0 (Katz et al. 2010), UCSC knownGene (Meyer et al. 2013), and Ensembl 71 (Flicek et al. 2013) annotations for the UCSC hg19 (NCBI GRCh37) human genome assembly, and a splice junction annotation file was created by enumerating all possible combinations of annotated splice sites as previously described (Hubert et al. 2013). RSEM (Li and Dewey 2011) and Bowtie (Langmead et al. 2009) were used to map reads to the gene annotation file, and TopHat (Trapnell et al. 2009) was used to align remaining unaligned reads to the genome and splice junctions (full details in Supplemental Methods).

### Isoform expression measurements

MISO (Katz et al. 2010) and v2.0 of its annotations were used to quantify isoform ratios for annotated alternative splicing events, and alternative splicing of constitutive junctions and retention of constitutive introns was quantified with junction reads as previously described (Hubert et al. 2013). All analyses were restricted to splicing events with at least 20 relevant reads (reads supporting either or both isoforms) that were alternatively spliced in our data. Events were defined as differentially spliced between two samples if they satisfied the following criteria: (1) at least 20 relevant reads in both samples, (2) a change in isoform ratio of at least 10%, and (3) a Bayes factor greater than or equal to 2.5 (AML data) or 5 (K562 data). Because the AML data had approximately two-fold lower read coverage than the K562 data, we reduced the Bayes factor by a factor of two to compensate for the loss in statistical power. Wagenmakers’s framework (Wagenmakers et al. 2010) was used to compute Bayes factors for differences in isoform ratios between samples. A description of the isoform-specific PCR used for Supplemental Figure S5 is given in Supplemental Methods. To identify splicing events that were differentially spliced in AML S34 samples vs. WT samples (Supplemental File S1), we used the Mann-Whitney U test and required *p* < 0.01. To identify splicing events that were differentially spliced in each AML sample with a *U2AF1* mutation (Fig. 4A, Supplemental Fig. S6), each *U2AF1* mutant sample was compared to an average *U2AF1* WT sample. The average *U2AF1* WT sample was created by averaging isoform ratios over all 162 *U2AF1* WT samples.

### Cluster analysis, sequence logos, and gene ontology enrichment

To perform the cluster analysis of AML transcriptomes (Fig. 1C) and K562 cells (Fig. 2C), we identified cassette exons that displayed changes in isoform ratios ≥10% across the samples, and then further restricted to cassette exons with at least 100 informative reads across all samples. An informative read is defined as a RNA-seq read that supports either isoform, but not both. We created a similarity matrix using the Pearson correlation computed from the z-score normalized cassette exon inclusion values, and clustered the samples using Ward’s method. Sequence logos were created with v1.26.0 of the seqLogo package in Bioconductor (Gentleman et al. 2004). Gene ontology analysis was performed with GOseq (Young et al. 2010), and is described further in Supplemental Methods.

### Western blotting

Protein lysates from K562 cells pellets were generated by resuspension in RIPA buffer and protease inhibitor along with sonication. Protein concentrations were determined using the Bradford protein assay. 10 ug of protein was then subjected to SDS-PAGE and subsequently transferred to nitrocellulose membranes. Membranes were blocked with 5% milk in Tris-buffered saline (TBS) for 1 hour at room temperature and then incubated with primary antibody 1:1000 anti-U2AF1 (Bethyl Laboratories, catalog no. A302-080A), anti-FLAG (Thermo, catalog no. MA1-91878), anti-Histone H3 (Abcam, catalog no. ab1791), or anti-alpha-tubulin (Sigma, catalog no. T9026) for 1 hour at room temperature. Blots were washed with TBS containing 0.005% Tween 20 and then incubated with the appropriate secondary antibody for 1 hour at room temperature.

### Minigenes

An insert containing the *ATR* genomic locus (chr3:142168344-142172070) or *EPB49* genomic locus (chr8:21938036-21938724) was cloned into the EcoRV site of pUB6/V5-HisA vector (Invitrogen) by Gibson assembly cloning (NEB). Site-directed mutagenesis was used to generate different nucleotides at the −3 position at 3′ splice site. Details of minigene transfection and real-time PCR specified in Supplemental Methods.

### *In Vitro* splicing

A pre-mRNA substrate transcribed from the AdML derivative HMS388 was used in all splicing reactions (Jurica et al. 2002; Reichert et al. 2002). T7 run-off transcription was used to generate G(5’)ppp(5’)G-capped radiolabeled pre-mRNA using UTP [α-^32^P] and K562 nuclear extracts were isolated following a published protocol (Folco et al. 2012) with a minor modification. Pre-mRNA substrates were incubated in standard splicing conditions and RNA species were separated in a 12% denaturing polyacrylamide gel and visualized using a phosphoimager (full details in Supplemental Methods). For quantification in Figure 5, each species was normalized by subtracting the background and then dividing by the number of uracil nucleotides in that species. The percentage of the second step products was calculated by dividing the second step species (spliced mRNA and lariat intron) by the total of all species in the lane.

### Protein structure prediction

Models of U2AF1 (residues 9-174) in complex with a RNA fragment extending from the 3′ splice site positions -4 to +3 were built by combining template-based modeling, fragment assembly methods, and all-atom refinement. Models were built using the software package Rosetta (Leaver-Fay et al. 2011) with template coordinate data taken from the UHM:ULM complex structure (Kielkopf et al. 2001) (PDB ID 1jmt: residues A/46-143) and the TIS11d:RNA complex structure (Hudson et al. 2004) (PDB ID 1rgo: U2AF1 residues 16−37 mapped to A/195-216; residues 155-174 mapped to A/159-179; RNA positions -4 to -1 mapped to D/1-4; RNA positions +1 to +3 mapped to D/7-9). The remainder of the modeled region (residues 9-15, 38-45, and 144-154) was built using fragment assembly (with templated regions held internally fixed) in a low-resolution representation (backbone heavy atoms and side chain centroids) and force field. The fragment assembly simulation consisted of 6000 fragment-replacement trials, for which fragments of size 6 (trials 1-3000), 3 (trials 3001-5000), and 1 (trials 5001-6000) were used. The RNA was modeled in two pieces, one anchored in the N-terminal zinc finger and the other in the C-terminal zinc finger, with docking geometries taken from the TIS11d:RNA complex. A pseudo-energy term favoring chain closure was added to the potential function to reward closure of the chain break between the RNA fragments. The fragment assembly simulation was followed by all-atom refinement during which all side chains as well as the non-templated protein backbone and the RNA were flexible. Roughly 100,000 independent model building simulations were conducted, each with a different random number seed and using a randomly selected member of the 1rgo NMR ensemble as a template. Low-energy final models were clustered to identify frequently sampled conformations (the model depicted in Figure 5A was the center of the largest cluster). We explored a range of possible alignments of the splice site RNA within the complex, with the final model selected on the basis of all-atom energies, RNA chain closure, manual inspection, and known sequence features of the 3′ splice site motif.

## DATA ACCESS

The RNA-seq data from K562 cells have been submitted to the NCBI Gene Expression Omnibus (GEO; http://www.ncbi.nlm.nih.gov/geo/) under accession number GSE58871.

## ACKNOWLEDGEMENTS

We thank Beverly Torok-Storb for project assistance and advice, and Sue Biggins, Toshi Tsukiyama, and members of the Bradley lab for comments on the manuscript. This research was supported by the Hartwell Innovation Fund (RKB, AR), Damon Runyon Cancer Research Foundation DFS 04-12 (RKB), Ellison Medical Foundation AG-NS-1030-13 (RKB), NIH/NCI P30 CA015704 recruitment support (RKB), Fred Hutchinson Cancer Research Center institutional funds (RKB), NIH/NCI training grant T32 CA009657 (JOI), NIH/NIDDK P30 DK056465 pilot study (JOI), NIH/NHLBI U01 HL099993 (AR), NIH/NIDDK K08 DK082783 (AR), the J.P. McCarthy Foundation (AR), the Storb Foundation (AR), and NIH/NIGMS R01 GM088277 (PB).

## AUTHOR CONTRIBUTIONS

JOI designed the molecular genetics and biochemistry experiments. AR designed the cell culture and *U2AF1* expression strategies. JOI, AR, BH, MEM, and ASZ performed experimental work, including cloning, cell culture, and flow cytometry. RKB and PB performed computational analyses and wrote the manuscript, with contributions from other authors. RKB and AR initiated the study.

## DISCLOSURE DECLARATION

The authors declare that no competing interests exist.

## Supplemental Material

**Supplemental Figure S1. Mutant *U2AF1* Expression Does Not Cause Splicing Failure.** (A) Levels of exon inclusion for NMD-irrelevant cassette exons in K562 cells expressing WT or mutant *U2AF1*, or transfected with a control siRNA or siRNA pool against *U2AF1*. (B) Levels of NMD-inducing isoforms of cassette exon events in K562 cells expressing WT or mutant *U2AF1*, or transfected with a control siRNA or siRNA pool against *U2AF1*. (C) Levels of properly spliced constitutive introns in K562 cells expressing WT or mutant *U2AF1*, or transfected with a control siRNA or siRNA pool against *U2AF1*.

**Supplemental Figure S2. *U2AF1* Mutations are not Associated with Increased Levels of NMD Substrates in AML Transcriptomes.** Plot illustrates the relative levels of NMD-inducing isoforms of alternatively spliced cassette exon events in each AML sample, with samples ordered by increasing level of NMD substrates. For each sample, all NMD-inducing isoforms that were increased or decreased ≥10% relative to the median over all samples were identified, and the quantity 100 × (# increased - # decreased) / (# increased + # decreased) was plotted on the vertical axis. Therefore, a positive value indicates a global increase in levels of NMD substrates, and vice versa. Samples with *U2AF1* mutations (black) do not exhibit higher levels of NMD substrates than do samples without *U2AF1* mutations (gray). Numbers above bars indicate the number of differentially expressed events for each sample.

**Supplemental Figure S3. *U2AF1* Mutations are not Associated with Increased Retention of Constitutive Introns in AML Transcriptomes.** Plot illustrates the relative levels of properly spliced out constitutive introns in each AML sample, with samples ordered by increasing level of proper splicing (intron removal). For each sample, all constitutive introns with evidence of increases/decreases in splicing ≥10% relative to the median over all samples were identified, and the quantity 100 × (# increased - # decreased) / (# increased + # decreased) was plotted on the vertical axis. Therefore, a positive value indicates a global increase in properly spliced constitutive introns, and vice versa. While retention of constitutive introns is common in AML transcriptomes—most bars are below 0—samples with *U2AF1* mutations (black) do not exhibit higher levels of constitutive intron retention than do samples without U2AF1 mutations (gray). Numbers above bars indicate the number of differentially retained constitutive introns for each sample; the plot is restricted to these events (e.g., the vast majority of constitutive introns are never retained in any sample, and those introns are not analyzed here since the plot is restricted to events that differ between samples).

**Supplemental Figure S4. *U2AF1* Mutations are not Associated with Increased Exon Skipping in AML Transcriptomes.** Plot illustrates the relative levels of inclusion of alternatively spliced cassette exons that are NMD-irrelevant in each AML sample, with samples ordered by increasing level of exon inclusion. For each sample, all NMD-irrelevant cassette exons whose inclusion was increased or decreased ≥10% relative to the median over all samples were identified, and the quantity 100 × (# increased - # decreased) / (# increased + # decreased) was plotted on the vertical axis. Therefore, a positive value indicates a global increase in cassette exon inclusion, and vice versa. Samples with *U2AF1* mutations (black) do not exhibit higher levels of cassette exon skipping than do samples without *U2AF1* mutations (gray). Numbers above bars indicate the number of differentially expressed events for each sample; the plot is restricted to these events. NMD-irrelevant cassette exons are defined as events for which either both or neither of the inclusion and exclusion isoforms are NMD substrates. The plot is restricted to NMD-irrelevant events to distinguish exon inclusion from NMD.

**Supplemental Figure S5. *U2AF1* Mutations Affect Genes Involved in Disease-Relevant Cellular Processes.** (A-F) Isoform ratios of splicing events illustrated in Figure 3C-E, G-I. Here, isoform ratios were measured by real-time PCR using primers specific to each isoform. Error bars, standard deviation estimated from five biological replicates. Panels A-F correspond to Figure 3C-E, G-I, in that order.

**Supplemental Figure S6. *U2AF1* Mutations are Associated with Altered 3′ Splice Site Consensus Sequences in AML Transcriptomes.** As Figure 4A, but for all seven AML samples with *U2AF1* mutations.

**Supplemental Figure S7. *U2AF1* WT AML Transcriptomes Do Not Exhibit Altered 3′ Splice Site Consensus Sequences.** As Figure 4A, but for seven randomly chosen AML samples without *U2AF1* mutations.

**Supplemental File S1. Differentially Spliced Events in AML Samples with S34 Mutations.** Splicing events that are differentially spliced in AML samples with S34F/Y mutations versus WT samples. Each row of the table corresponds to isoform 1 of a splicing event, where isoform 1 is defined as follows: inclusion isoform for cassette exons (“se”), most intron-proximal isoform for competing 5′ and 3′ splice sites (“a5ss”, “a3ss”), inclusion of upstream exon for mutually exclusive exons (“mxe”), splicing of retained introns annotated as alternative (“ri”) or constitutive (“ci”), and canonical splicing of constitutive junction (“cj”). Each row is assigned a unique identifier specifying the event type and coordinates of the upstream junctions for isoforms 1 and 2 of the event; this event identifier format is a modification of the format used by MISO (Katz et al. 2010). The columns of the table are defined as follows: “coords”, genomic coordinates containing the event; “spliceSites”, dinucleotides at the 5′ and 3′ splice sites of the upstream junction of isoform 1; “nmdTarget”, whether the specified isoform is a predicted NMD target, where a value of “NA” indicates that the event is not NMD relevant (e.g., neither or both isoforms are predicted substrates for NMD); “deltaPsi”, difference in isoform ratio between the two sample groups; “pval”, *p*-value computed with the Mann-Whitney U test for group comparisons; “gene”, gene ID; “geneName”, gene name; “geneDescription”, gene description. Gene IDs, names, and descriptions are from Ensembl, when available.

**Supplemental File S2. Differentially Spliced Events in K562 Cells Expressing S34F.** As Supplemental File S1, but for K562 cells expressing S34F versus WT *U2AF1*. Here, the “deltaPsi” column specifies the difference in isoform ratio between the mutant vs. WT cells, and “pval” is replaced by “bayesFactor”, the Bayes factor associated with the sample comparison.

**Supplemental File S3. Differentially Spliced Events in K562 Cells Expressing S34Y.** As Supplemental File S2, but for S34Y expression.

**Supplemental File S4. Differentially Spliced Events in K562 Cells Expressing Q157P.** As Supplemental File S2, but for Q157P expression.

**Supplemental File S5. Differentially Spliced Events in K562 Cells Expressing Q157R.** As Supplemental File S2, but for Q157R expression.

**Supplemental File S6. Theoretical Model of the U2AF1:RNA Complex.** Computationally predicted model encompassing 3′ splice site residues -4 to +3 built using fragment assembly. Multi-model PDB file contains the center (model 1) and 19 randomly selected members (models 2-20) of the largest cluster after structure-based comparison and clustering of all low-energy models.

